# OCHROdb: a comprehensive, quality checked database of open chromatin regions from sequencing data

**DOI:** 10.1101/484840

**Authors:** Parisa Shooshtari, Samantha Feng, Viswateja Nelakuditi, Justin Foong, Michael Brudno, Chris Cotsapas

## Abstract

International consortia, including ENCODE, Roadmap Epigenomics, Genomics of Gene Regulation and Blueprint Epigenome have made large-scale datasets of open chromatin regions publicly available. While these datasets are extremely useful for studying mechanisms of gene regulation in disease and cell development, they only identify open chromatin regions in individual samples. A uniform comparison of accessibility of the same regulatory sites across multiple samples is necessary to correlate open chromatin accessibility and expression of target genes across matched cell types. Additionally, although replicate samples are available for majority of cell types, a comprehensive replication-based quality checking of individual regulatory sites is still lacking. We have integrated 828 DNase-I hypersensitive sequencing samples, which we have uniformly processed and then clustered their regulatory regions across all samples. We checked the quality of open-chromatin regions using our replication test. This has resulted in a comprehensive, quality-checked **d**ata**b**ase of **O**pen **CHRO**matin (**OCHROdb**) regions for 194 unique human cell types and cell lines which can serve as a reference for gene regulatory studies involving open chromatin. We have made this resource publicly available: users can download the whole database, or query it for their genomic regions of interest and visualize the results in an interactive genome browser.

## INTRODUCTION

In recent years, multiple international consortia-based projects, such as ENCODE (1), NIH Roadmap Epigenomics Mapping Consortium (REMC) Project (2), NHGRI Genomics of Gene Regulation (GGR) Project (3) and Blueprint Epigenome (4) have made several large scale chromatin accessibility datasets publicly available. ENCODE, GGR and REMC datasets contain open chromatin data samples (e.g. ATAC-seq and DNase-I seq) for hundreds of primary cell types, stem cells, cell lines and tissues. Blueprint Epigenome project has also made open chromatin data available for hundreds of haematopoietic cell types. These datasets are extremely useful for studying mechanisms of gene regulation in different contexts, including disease and cell development. However, current regulatory region identification pipelines process samples within these datasets individually, making it difficult to annotate and analyze DHS sites across multiple samples in a study, and across studies. Accessibility of regions of open chromatin varies across cell types, and may even vary within biological replicates of the same cell type. The absence of a well-defined, reference map for genomic locations of accessible chromatin that is also consistent across replicates frequently poses challenges to many downstream analyses, such as investigations into the association between accessibility state of specific regulatory elements and the expression of the genes which they control.

Although most epigenomics datasets such as ENCODE, GGR, REMC and Blueprint Epigenome data contain multiple biological and/or technical replicates of the same cell type, a comprehensive replication-based quality check of individual regulatory regions identified by peak calling algorithms is still lacking. Quality checking measurements that are used in other open chromatin databases check quality of individual samples and filter unreliable samples (5); however, they do not assess quality of individual regulatory sites across multiple samples. It is already well-recognized that several peaks identified in a sample by peak calling algorithms may not be supported in replicate samples (6) (7). Quality checking methods such as jMOSAiCS (6), MSPC (7) and irreproducibility discovery rate (IDR) (8) have been developed to compare peaks identified in pairs of replicates in order to identify replicable peaks in sequencing experiments with a main focus on ChIP-seq experiments. IDR in particularly has been used in the ENCODE Project, and has been applied to various sequencing-based experiments such as ATAC-seq, DNase-seq and ChIP-seq data. However, QC methods are limited by the fact that they compare replicates from one cell type only, and subsequently can be quite stringent at filtering true peaks in the absence of orthogonal information.

In order to address the above problems we designed and implemented a pipeline that obtained DNase-I Hypersensitive (DHS) peaks of 828 data samples obtained from multiple resources (including ENCODE, REMC, Blueprint Epigenome and GGR), aligned them across samples, assessed replicability of each DHS site across several cell types using our recently developed replication test (9) and filters unreliable DHS sites (9). This has resulted in a quality-checked database of regions of open chromatin comprising 1,455,046 reliable DHS for 194 unique cell types across the autosomal genome. To our knowledge, OCHROdb is the first and the most comprehensive DHS database obtained by (a) aligning the same regulatory sites across more than 800 samples from multiple public datasets and (b) checking quality of individual DHS sites, and it can serve as a reference for studies concerned with DHS activities and their role in gene regulation. This database can be accessed through our website (https://dhs.ccm.sickkids.ca/), where users can query and download the data either fully or at specific genomic regions of interest. We have also prepared an interactive genome browser for effective visualization of DHS sites across several cell types.

## MATERIAL AND METHODS

### Data collection

We collected DHS data generated by multiple international consortia-based projects including ENCODE, REMC, GGR and Blueprint Epigenome. The data generated by ENCODE, REMC and GGR are hosted by ENCODE and publicly available through (https://www.encodeproject.org). Blueprint DHS data samples from Haematopoietic Epigenomes are available at (http://www.blueprint-epigenome.eu). In August 2017 we collected from these resources the following types of the data:

i. Data samples in BED format, where regions of open chromatin were identified by Hotspot peak calling algorithm (narrow peaks; FDR of 0.05) (10). This was available for 362 samples from ENCODE, 318 samples from REMC, 51 samples from GGR and 97 samples from Blueprint Epigenome project (See Table S1 for a full list of samples).
ii. Metadata describing file accession ID, cell type, project, assembly and file download URL (See metadata in Tables S1). We used ENCODExplorer R Bioconductor package (11) to get access to and query metadata for ENCODE, REMC and GGR data files. The metadata of Blueprint Epigenome data was downloaded from ftp://ftp.ebi.ac.uk/pub/databases/blueprint/data_index/homo_sapiens/data.index.

### Data processing workflow

#### Step 1 – Analyzing raw DNase-I sequencing data to identify DHS peaks

All DHS samples released from ENCODE project website (i.e. GGR, ENCODE and REMC data samples) had previously gone through the ENCODE uniform processing pipeline for DNase-I Hypersensitive experiments (https://github.com/ENCODE-DCC/dnase_pipeline/). Briefly, first a bwa index and a mappability file were produced by using a reference (e.g. GCRh38) fasta file and a DNase mappability file specific to a read size. Then BWA was used to generate bam files containing all reads mapped to the reference genome, and the bam files were merged and filtered to include only high quality mapped reads. At the last step, hotspot peak calling algorithm (10) was applied to the filtered bam files to identify enriched regions (i.e. peaks and hotspots). DHS data samples released through Blueprint Epigenome project were also analyzed following a similar procedure. First, the reads were mapped to human genome GCRh38 reference using BWA 0.7.7. Bam files were sorted and duplicates were marked using Picard Tools (12). Then, the output bam files were filtered and reads with Mapping Quality of less than 15 were removed. Finally, the Hotspot peak calling algorithm was applied to identify peaks and hotspots.

#### Step 2 – Checking quality of samples

We originally obtained a total of 847 samples containing DHS peaks in BED format from ENCODE and Blueprint site (Figure 1A). We found that five Blueprint data samples do not have a cell type and/or donor ID assigned to them in the metadata and removed them. Additionally, by considering distribution of number of DHS peaks per sample, we identified and removed 14 outlying samples with too many (> 500,000 peaks) or too few (< 40,000 peaks) DHS peaks (See Figure 3A for distribution of number of DHS peaks per samples). The total number of samples remaining after the above quality checking steps was 828 (Figure 1A). Of these 828 samples, 45 were mapped to GRCh37 genome reference and the rest were mapped to GRCh38 genome build. We converted GRCh37 samples to GRCh38 using the GATK API (13). Figure S1 shows a histogram of the number of biological and technical replicates per cell type. As shown in this figure, these 828 samples come from 194 unique cell types. 161/194 cell types have at least two replicate samples.

**Figure 1:**
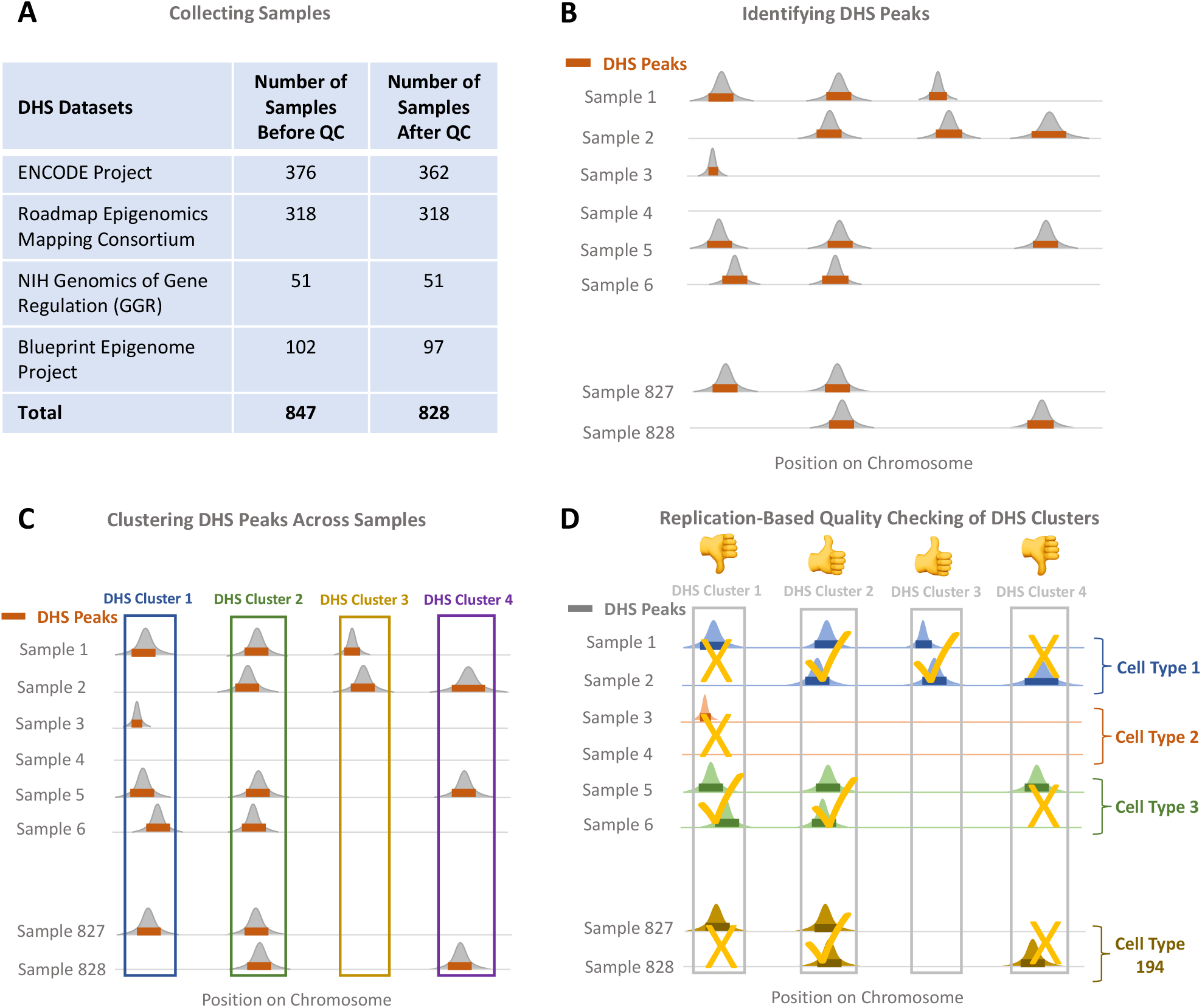
Post processing of DNAse-I Hypersensitive (DHS) Data. (A) A total of 847 DHS data samples were collected from ENCODE, REMC, GGR and Blueprint Projects. 828/847 passed initial quality checking. (B) All the DNase-I sequencing data samples were already pre-processed by ENCODE and Blueprint consortia and narrow peaks were identified by Hotspot peak calling algorithm (10). (C) Following the method we previously developed (9), we clustered DHS peaks across samples and (D) checked quality of each DHS cluster using our replication test.

#### Step 3 – Clustering peaks

In the 828 samples, we found a total of 117,488,197 DHS peaks on the autosomes (i.e. excluding X and Y chromosomes). Peak calling algorithms such as Hotspot (10) and MACS (14) identify peaks in each individual sample separately (Figure 1B). In order to identify the same DHS sites over multiple samples, we followed the method that we developed earlier (9), where we used Markov clustering (MCL) (15) with its default parameters to cluster the peaks across multiple samples (Figure 1C). In our application, each DHS peak is treated as a graph node. Similarity between the two nodes is defined as the length of intersection between the two peaks divided by the length of union of the intervals containing of the two peaks. Although any other clustering algorithm can be used at this stage, we chose MCL for two reasons. First, MCL does not require estimation of the number of clusters *a priori*. Second, the complexity of MCL is *O*(*Nk*^2^), where *N* is the number of nodes (i.e. peaks), and *k* is the number of resources allocated per node (a relatively small number for sparse graphs). The fact that MCL complexity is linear with the number of peaks makes it quite fast for our application. Using MCL we were able to group 117,488,197 DHS peaks across 828 samples into 4,020,940 DHS clusters across the autosomal chromosomes.

#### Step 4 – Replication-based quality checking of DHS Clusters

Both peak calling and peak clustering ignore sample labels (i.e. cell types of origin), so we can check DHS cluster quality by looking for evidence that DHS peaks are replicated in this analysis. We expect that for a DHS cluster that represents a true regulatory site, the DHS accessibility is consistent across replicates. We therefore employed our previously designed replication test (9) to assess consistency across replicates (Figure 1D). As required by our replication test, we selected cell types with at least two replicates (161/194 of cell types). Since for 84/161 cell types, we had more than two replicates per cell type (Figure S1), we repeated our replication test ten times, each time we randomly selected two replicates per cell type. We then merged the test statistics obtained from each test. The output of each run of our replication test follows a Chi-squared distribution with one degree of freedom (*χ*^2^ (df=1)) (9) and thus, the sum of the test statistics resulting from running our replication test ten times follows a *χ*^2^ (df= 10) distribution (16). We selected DHS clusters that pass a nominal significance threshold of p ≤ 0.05 for the combined replication test and called them replicable DHS. In order to examine how many runs of replication test is required in order to obtain a stable set of replicable DHS, we applied the replication test 20 times. Then for any N between 1 and 19, we found a list of replication test obtained by combining N replication tests. Our results indicated that the Area Under the Curve (AUC) obtained for sensitivity/specificity analysis remained above 0.95 after nine iterations, confirming that running the replication test ten times is appropriate (Figure S2). 1,460,986/4,020,940 (36.3%) of DHS clusters passed the combined replication test. As we previously showed, active replicable DHSs that we annotated capture the majority of proportion of disease heritability (h2g) explained by all DHS-detected peaks in a tissue (9), suggesting the non-replicating peaks are spurious.

#### Step 5 – Incorporating DHS accessibility significance

Chromatin accessibility as a quantitative measure is the density of mapped DNase-I cleavage at different genomic locations. Chromatin accessibility varies across replicable DHS sites of the same sample. Also, different samples have different accessibilities at the same replicable DHS site. Since DHS accessibility can be an indicator of how active a DHS site is in a particular sample, we incorporated DHS accessibility information to our data. To measure significance of accessibility of each replicable DHS in each sample, we first identified all DHS peaks that belong to a replicable DHS, and assigned *-log10*(*P value*) of the most significant DHS peak of a sample to that replicable DHS in the sample of interest.

There exist batch effects in DHS intensities (i.e. -*log10*(*P value of accessibility*)) due to the fact that DHS data were generated and processed in multiple centres (ENCODE, REMC, Blueprint and GGR) (Figure 2A). As the result of this batch effect, in the tSNE plot, samples generated by the same project grouped together disregard of their cell type similarity (Figure 2B). We applied the following steps to remove these batch effects. First, we normalized DHS intensities within each sample by linearly scaling them between 0 and 1. The number of accessible DHS varied from sample to sample. We therefore randomly selected 10,000 accessible DHS sites (i.e. with non-zero intensities) from each sample to estimate DHS intensity distributions for each sample, and adopted a method similar to ComBat (17) (18) to remove batch effects. Briefly, we sorted intensities of randomly selected DHS sites within each sample, quantile normalized them to make distributions of intensities similar across samples. We then interpolated intensities of the DHS sites that were not initially chosen in the random selection and estimated their quantile normalized values. This process resulted in removing batch effects of DHS intensities, and made them comparable across samples generated by different centres, and therefore samples grouped together based on their cell type similarity (Figure 2C).

**Figure 2:**
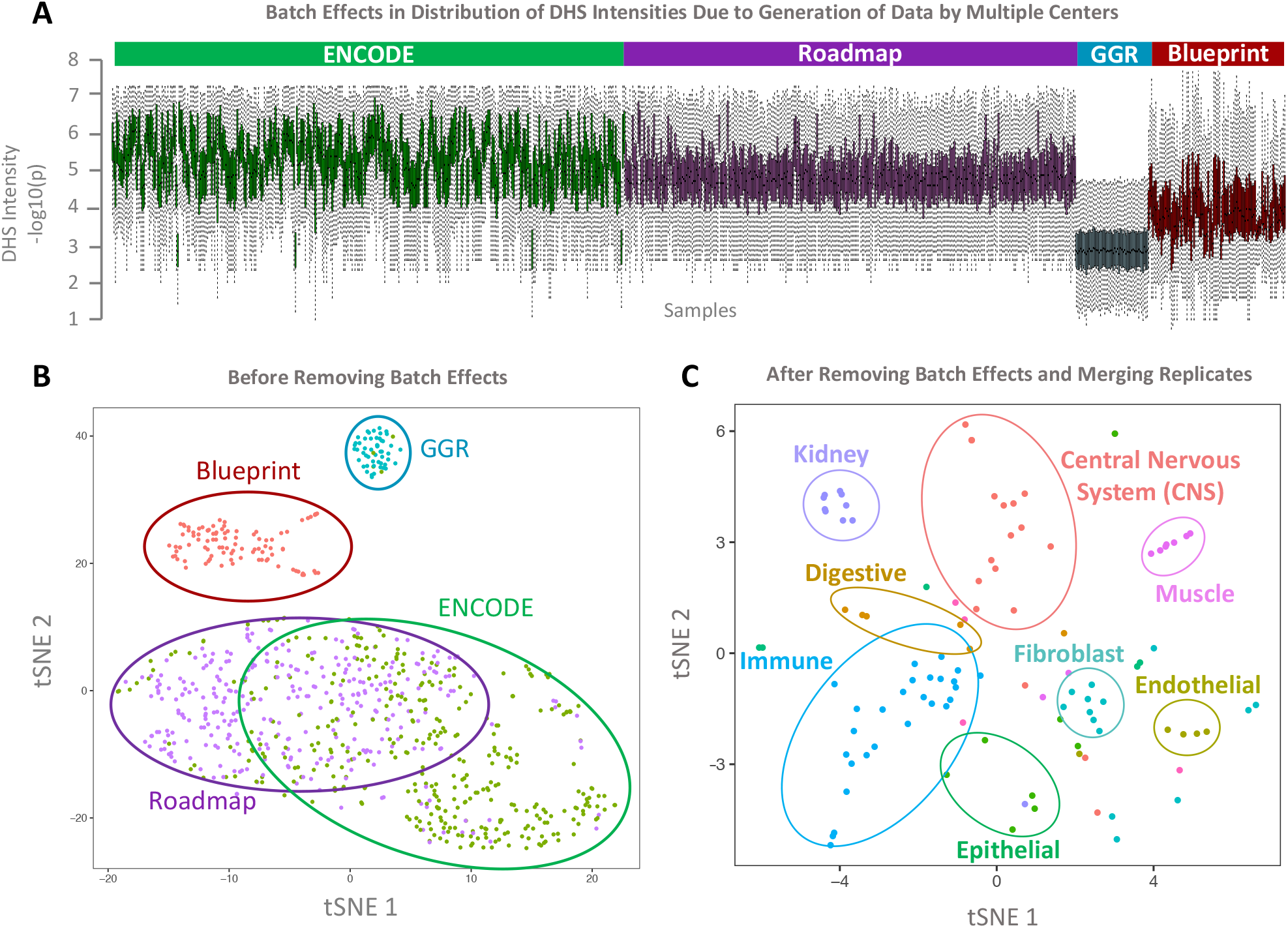
Removing batch effects caused by generation of data by multiple centers. (A) There exist batch effects in accessibility levels of replicable DHS clusters as a result of generation of data by multiple centers. (B) This resulted in grouping samples together based on the project that generated them disrespect of cell type similarity. (C) We removed batch effects and merge intensities of multiple replicates of the same cell types. After batch effect removal, in tSNE plot cell types grouped together based on their cell type similarities.

Finally, we collapsed DHS intensities of samples from the same cell types by measuring median intensities over these samples. 1,455,046 out of 1,460,986 (99.5%) replicable DHS have median intensities higher than 0.25 in at least one cell type. We considered these 1,455,046 sites as the final replicable DHS sites to build our database of regions of open chromatin for 194 cell types.

## RESULTS

### Database content and statistics

We have prepared a comprehensive and quality-checked database of regions of open chromatin using DNase-I Hypersensitive datasets generated by REMC, ENCODE, GGR and Blueprint Epigenome Projects. OCHROdb contains DHS intensities for 1,455,046 DHS sites aligned over 828 samples. These 828 samples comprise of 194 distinct cell types, including 27 immune cell types, 18 CNS, 9 muscle, 57 cell lines, 10 stem cells, 6 digestive system cell types in addition to lung, cardiovascular, fibroblasts, kidney, extra embryo, male and female reproductive system and endothelial cell types (See Table S1 for the full list of cell types). tSNE plots of DHS activities of the 194 cell types shows that cell types from similar tissues group together confirming they have similar DHS activities (Figure 2C).

As it is shown in Figure 3C, the number of accessible replicable DHS varies across cell types. Additionally, number of cell types where a replicable DHS is accessible can be any number between one cell type (i.e. cell-type specific DHS) to 194 cell types (i.e. constitutive DHS). The median number of active cell types per replicable DHS is eight (mean=21.3; sd=33.4) (Figure 3B). Around 10.2% (148,142/1,455,046) of replicable DHS are cell-type specific, meaning that they are active in one cell type, and only 1,773 out of 1,455,046 replicable DHS are constitutive, meaning that they are active in all 194 cell types. Median length of replicable DHS is 310 base pairs (sd = 112 bp) (Figure 3B).

**Figure 3:**
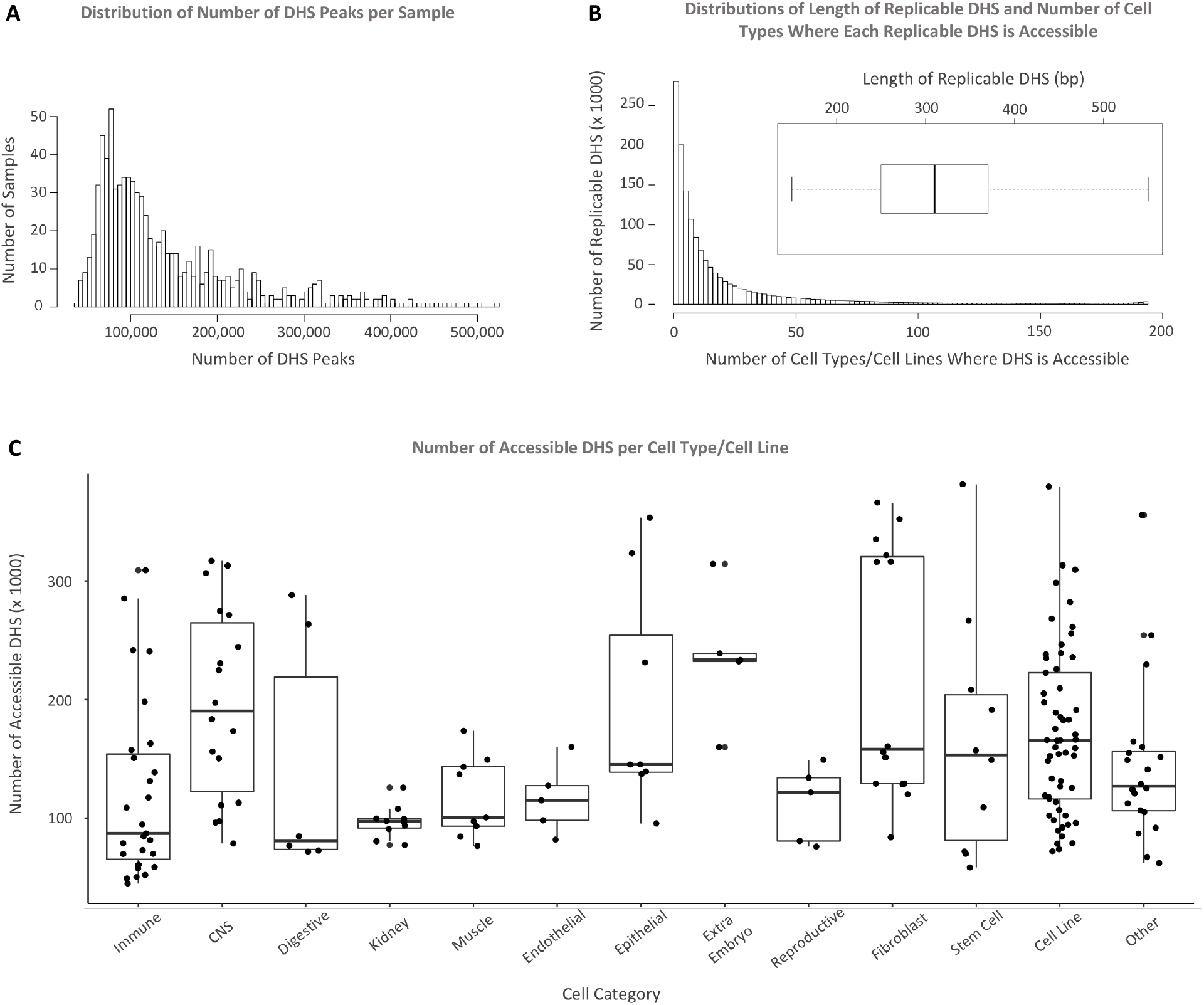
Statistics on DHS data. (A) Distribution of number of DHS peaks per sample. (B) Distributions of length of replicable DHS and number of cell types where each replicable DHS is accessible. Average length of replicable DHSs is 310 base pairs. Around 10.2% of replicable DHS are active in one cell type only and a very small portion of DHS (1,773/1,455,046) are active in all cell types. (C) Number of replicable DHS is between around 40,000 to 400,000, and it varies substantially across cell types. The percentage of genome covered by replicable DHS per cell type is between around 0.4% to 4% depending on the cell type.

### Web interface and genome browser

We developed a web interface that permits users and researchers access to OCHROdb by primarily utilizing React, Tabix, Node.js, and Express.js. React (19), a component-based JavaScript library developed by Facebook, was selected for the front-end to establish a single page application with high performance. React helps to render HTML for web applications without refreshing the page or website templates and allows the client and server to communicate faster. Tabix is a software package that indexes tab-delimited files to efficiently perform queries and was developed specifically for biological data. Data indexing involves sorting data based on specific fields and allows a query to be completed without reading the entire data file, greatly reducing query speeds if implemented properly. Several actions were executed to prepare the DHS dataset for Tabix queries, such as sorting the files by chromosome and start position, compressing the file, and indexing the file with Tabix. The back-end of our web application is composed of the server-side JavaScript runtime environment Node.js (20), in conjunction with Express.js (21), a web application framework that is used for the web server. Node.js forms a connection between Tabix (22) and React front-end and allows the user to view and interact with the DHS data on the web application. Express.js is a flexible and highly documented web framework built on top of Node.js and greatly simplifies the complexity of back-end code written.

We have made the metadata, entire curated DHS dataset and data specific to each chromosome downloadable from the web interface. Through the web interface, the user can also query the database by specifying a region of interest (i.e. entering a specific chromosome number and start and end coordinates). After submitting coordinates of a region of interest, an exportable table of the results matching the user input is generated by a JavaScript library called DataTables.net (23) (Figure S3A).

Additionally, the user can visualize the replicable DHS data through JBrowse (24), an embeddable genome browser, by specifying coordinates of the region and cell types of interest (Figure 4). To do this, a GFF (general feature format) file per cell type is generated by reading the original BED files using a Python script. Information specific to each replicable DHS (e.g. intensity of the DHS for any given cell-type and genomic coordinates of the DHS) can be found in the pop-up window that opens after clicking on a given DHS (Figure S3B). Replicable DHS are only visible if they are accessible (i.e. having non-zero intensities) at the cell types of interest. The replicable DHS that are accessible in multiple cell types get the same color on different tracks (each corresponding to a cell type) of the genome browser, to make adjacent DHS more trackable visually across several cell types.

**Figure 4:**
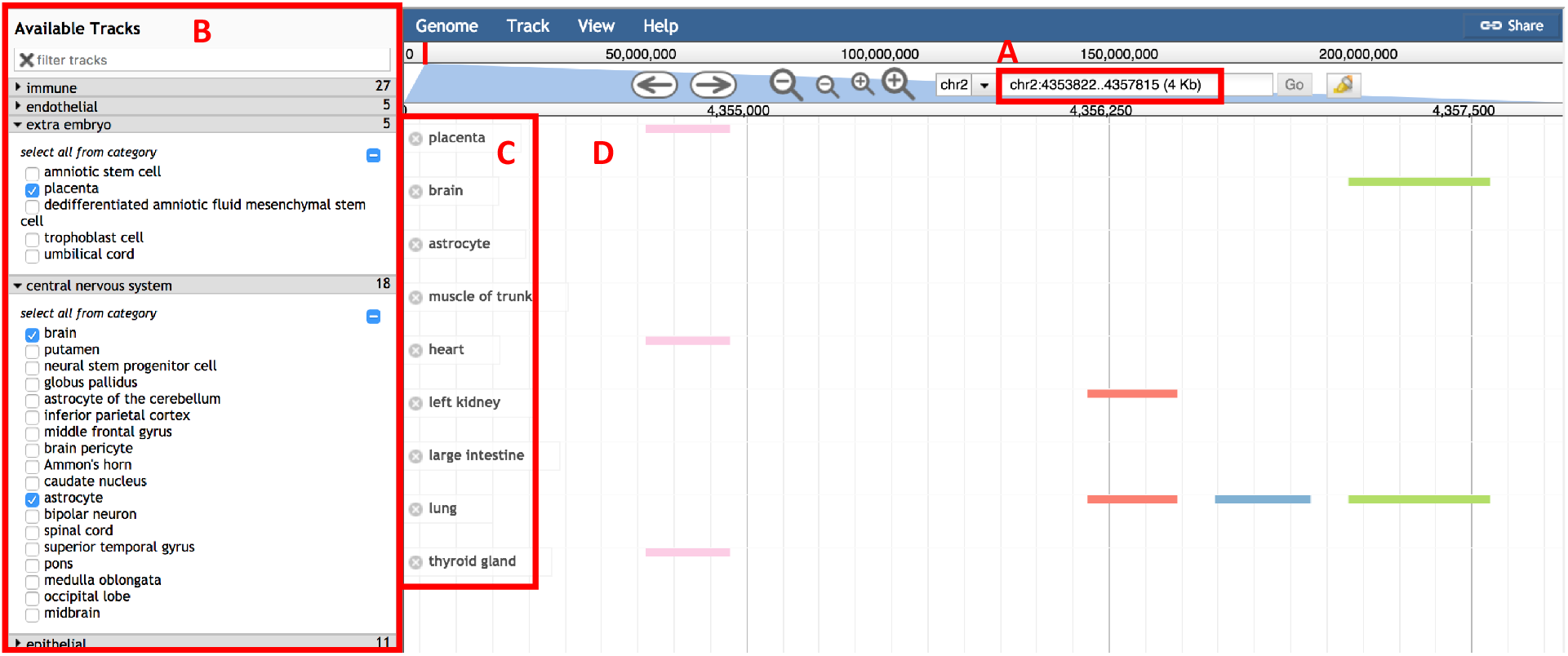
Interactive genome browser. We developed a customized genome browser to visualize DHS data. Here users can (A) specify their genomic region of interest, and (B) select from different cell categories and different cell types within each category. (C) The selected cell types appear on the left side of the browser. (D) Genomic location of replicable DHS appears on the middle of the browser, where each replicable DHS, aligned across multiple cell types, appears in a different colour to make it distinguishable from other replicable DHS adjacent to it.

## DISCUSSION

Multiple large-scale consortia-based projects, including ENCODE, REMC, Blueprint and GGR have generated thousands of sequencing-based data samples that capture regions of open chromatin for the whole genome in hundreds of cell types. Building on this immensely informative DHS datasets, we have developed an analysis pipeline that gets hundreds of pre-processed DHS data samples (i.e. in narrow peaks format) as the input, aligns regions of open chromatin across samples, checks quality of each region using a replication-based test, and outputs a database of open chromatin accessibility across the whole genome.

Through applying our processing pipeline to 828 DHS data samples obtained from multiple public datasets, we have built a database of 1,455,046 regions across the whole genome for 828 samples comprising of 194 cell types. The main advantages of OCHROdb compared to previous datasets that are available publicly (e.g. ENCODE, REMC, Blueprint and GGR) are as follows.

i. In the publicly available datasets the DHS peaks are identified for each individual sample separately, while our method incorporates peaks from multiple samples together, aligns them across samples, and releases them as a coherent DHS database consisting of replicable peaks from multiple samples. Since in the previous datasets, each sample is analyzed separately, annotation of the same DHS sites across several samples is not available.
ii. Additionally, the batch effects that exist across samples due to the fact that multiple centers generate and analyze the data are not adjusted in the previous DHS datasets. In our database, we removed the batch effects that exist in the data samples collected from multiple centers.
iii. We have employed our replication test that considers multiple samples from a diverse range of cell types in order to check replicability of each DHS site. This results in identifying replicable DHS that present true regulatory sites (See partitioning heritability results in (9)), while preserving regulatory sites filtered by more stringent tests such as IDR (8).

Although we have explained our processing pipeline in the context of DHS data samples, it can also be applied to the narrow peaks obtained by other assays such as ATAC-seq. We therefore plan to process open chromatin datasets generated by other assays (e.g. ATAC-seq) and make these results available in our database. Additionally, we aim to process non-human (e.g. mouse) open chromatin datasets and release them separately in our open chromatin database.

Recently, International Human Epigenome Consortium (IHEC) have collected and processed individual epigenome data samples from multiple large-scale international projects, such as ENCODE, REMC and Blueprint Epigenome. It is expected that more epigenome samples from other projects become available through IHEC portal. This includes the data from Deutsches Epigenome Program (DEEP), McGill Epigenomics Mapping Center, 4D Nucleome, Hong Kong Epigenomes Project (EpiHK), Multiple MS and SYSCID. As more open chromatin data samples are generated and released by IHEC and other projects (25), we plan to apply our processing pipeline to these samples and release the processed data as part of our open chromatin database. Making a comprehensive database of regions of open chromatin using the data released by IHEC project aligns with IHEC goals and will benefit the scientific community. We will maintain our database and release new updates every six months.

In summary, we believe that our open chromatin database, which contains a wide range of cell types can serve as a reference map for regions of open chromatin and will find many applications in studies concerned with uncovering gene regulatory mechanisms of disease and cell development.

## Supporting information

## AVAILABILITY

OCHROdb is a database of open chromatin regions from sequencing data. It is available to download through https://dhs.ccm.sickkids.ca/. Additionally, our interactive open chromatin browser is accessible through the same link.

## SUPPLEMENTARY DATA

Supplementary Data are available at NAR online.

## ACKNOWLEDGEMENT

The authors would like to acknowledge Jonah Librach and Arun Ramani for editorial comments, and High Performance Computing team at the Hospital for Sick Children (SickKids) for helping with the web server.

