## Supplementary material for "OCHROdb: a comprehensive, quality checked database of open chromatin regions from sequencing data"

### Supplementary Data (Three supplementary figures and one supplementary table)

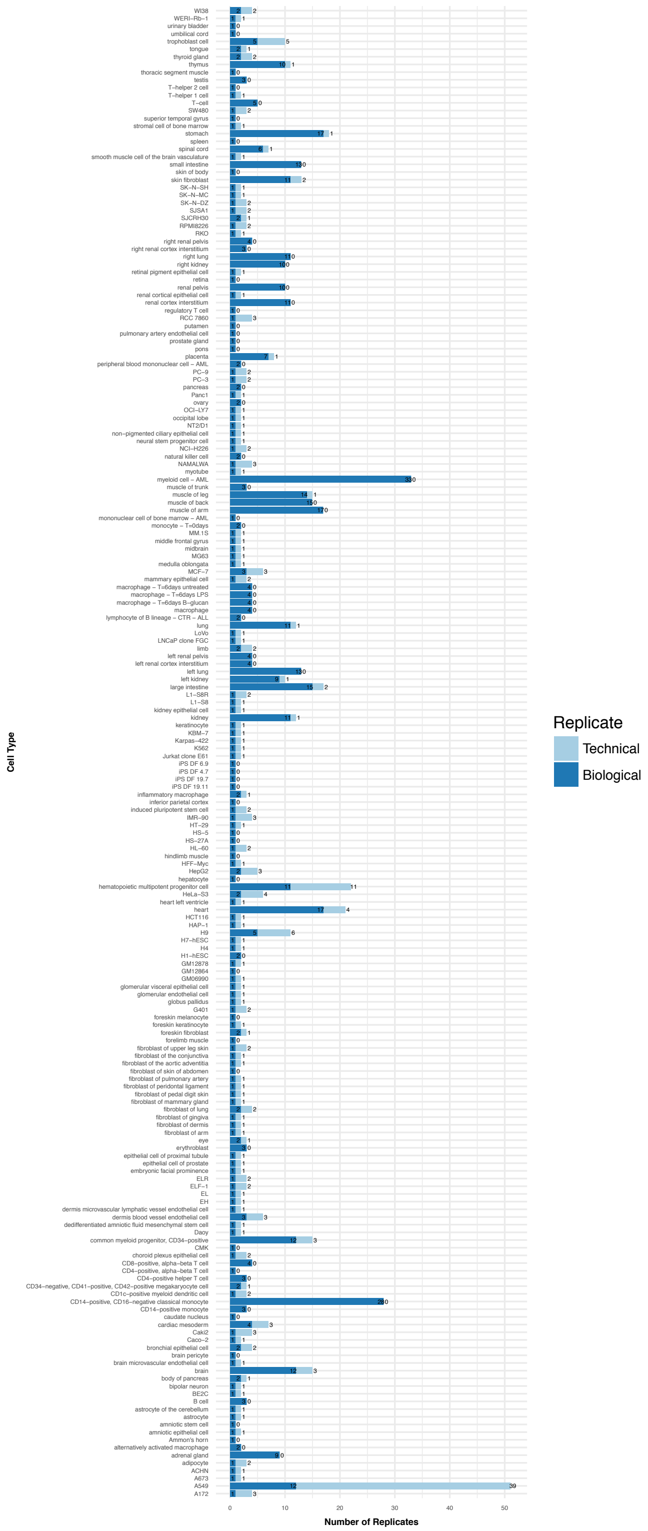


**Figure S1: Number of replicate samples per cell type.** For majority of cell types (161/194), there are at least two replicates per cell type. Number of technical and biological replicates varies across cells.

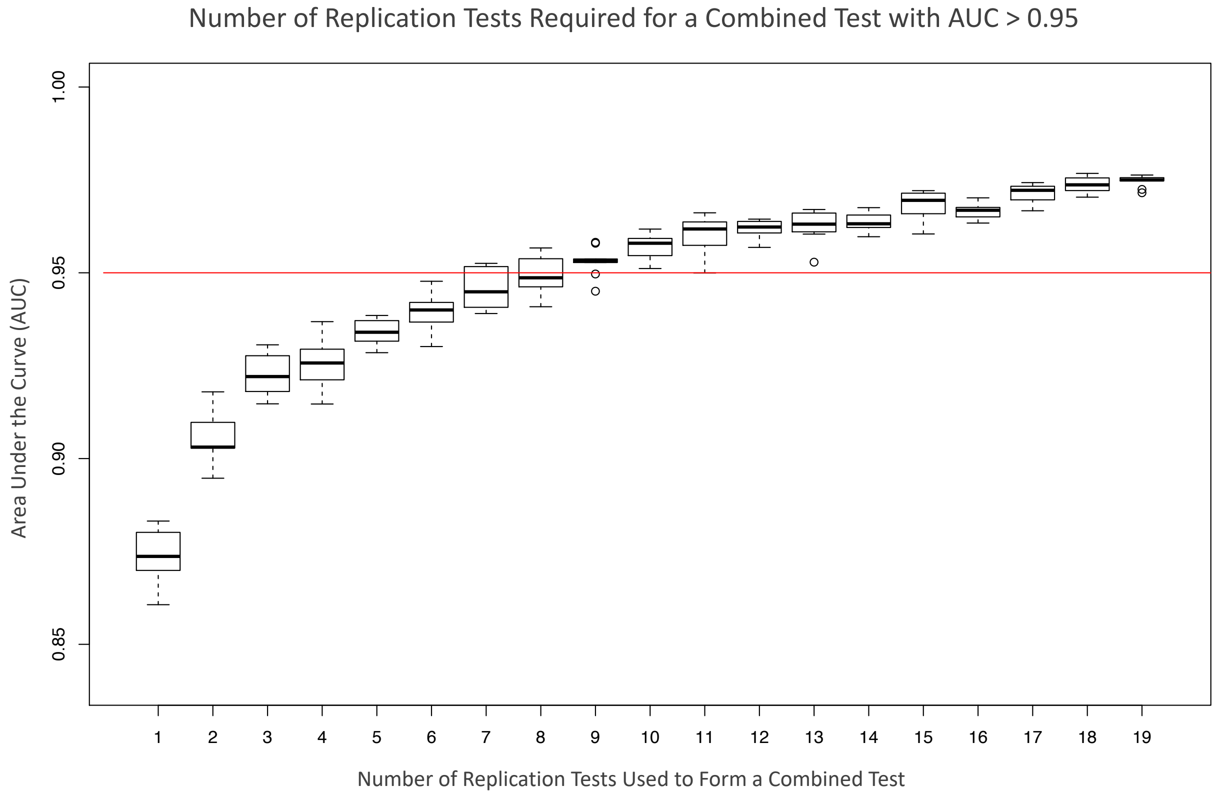


**Figure S2: Verifying number of replication tests required to accurately identifying replicable DHS.** We performed replication tests multiple times and combined them to identify replicable DHS clusters. Area under the curve remains above 0.95 after nine replication tests, highlighting that using ten replication tests is adequate for accurate identification of replicable DHS.

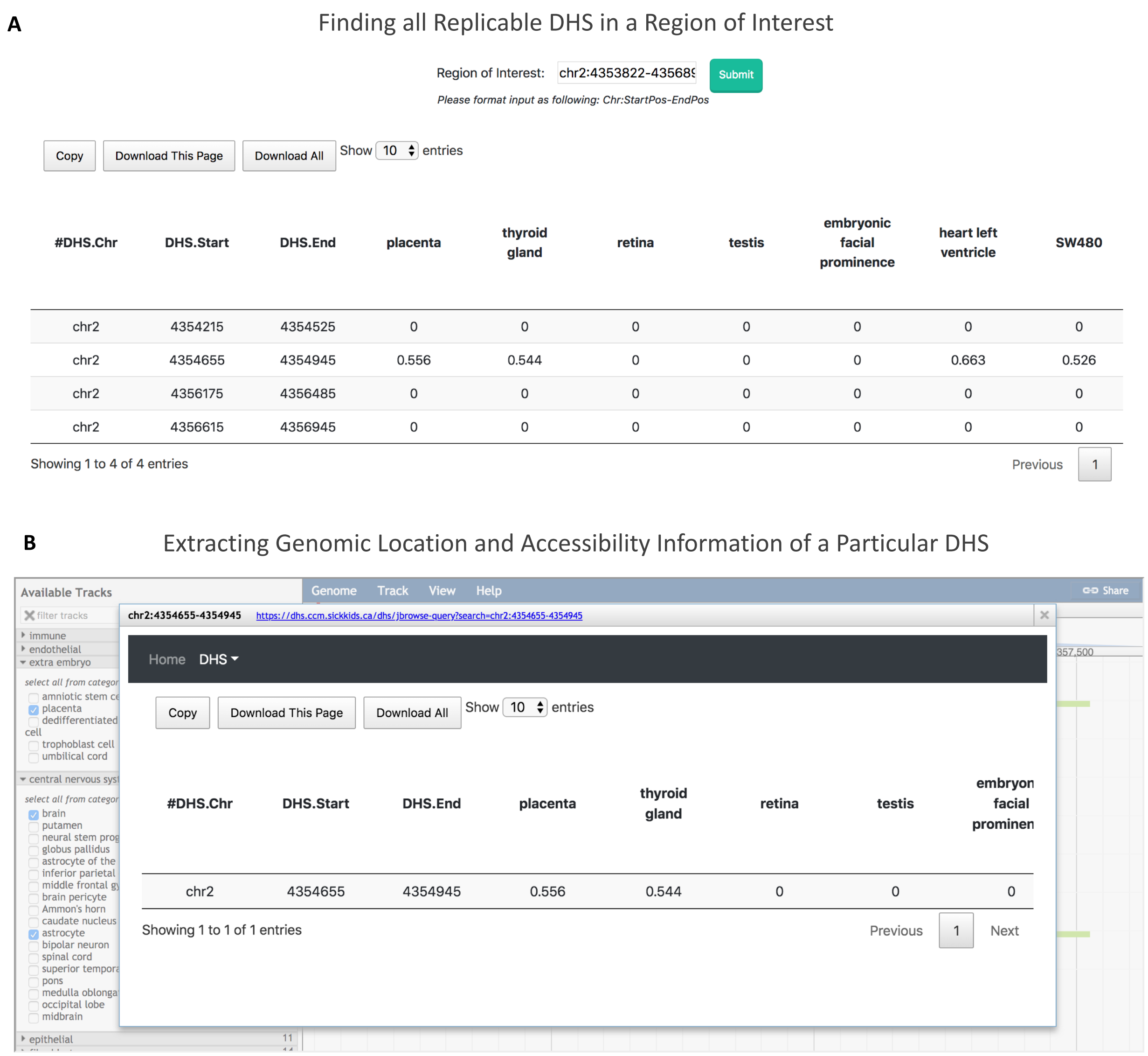


**Figure S3: Obtaining all replicable DHS overlapping a genomic region of interest and extracting information of a particular DHS.** Users can either download the full DHS data or (A) query their genomic region of interest and download region-specific replicable DHS. This will result in a file in Bed format containing genomic location and accessibility of each replicable DHS across 194 cell types. (B) In the interactive genome browser (Figure 4), user can click on a specific DHS and extract its information including its genomic location and accessibility levels across all cell types.

**Table S1: Information of 828 samples.** This table lists information of 828 samples, including the file accession ID in the ENCODE/Roadmap/GGR/Blueprint Project, biosample term name, biosample type, project, genome assembly and file download URL.
